# QuAPPro: An R/shiny app for Quantification and Alignment of Polysome Profiles

**DOI:** 10.1101/2024.05.02.592260

**Authors:** Chiara Schiller, Sonja Reitter, Janina A. Lehmann, Kai Fenzl, Johanna Schott

**Affiliations:** Mannheim Institute for Innate Immunoscience (MI3) and Mannheim Cancer Center (MCC), Medical Faculty Mannheim, Heidelberg University, Franz-Volhard-Str. 6, Mannheim, 68167, Germany; Center for Molecular Biology of Heidelberg University (ZMBH) and German Cancer Research Center (DKFZ), DKFZ-ZMBH Alliance, Im Neuenheimer Feld 329, Heidelberg, 69120, Germany; Institute for Computational Biomedicine, Faculty of Medicine, Heidelberg University Hospital and Heidelberg University, Im Neuenheimer Feld 130, Heidelberg, 69120, Germany; European Molecular Biology Laboratory (EMBL), Genome Biology Unit, Meyerhofstrasse 1, Heidelberg, 69117, Germany

**Keywords:** translation, protein biosynthesis, polysomes, polysome profiling, R shiny

## Abstract

Polysome profiling is a powerful technique to study mRNA translation. After separation of ribosomal subunits from monosomes and polysomes by ultracentrifugation on sucrose density gradients, a UV absorbance profile is recorded during elution. This profile can be used to assess global translational activity, or reveal changes in ribosome biogenesis or translation elongation. In parallel to UV absorbance profiles, it is also possible to record fluorescence to measure the association of fluorescently tagged proteins with ribosomes or polysomes. To this end, the area under subsections of the UV/fluorescence profiles needs to be quantified carefully. In addition, alignment of profiles in one graph helps to visualize differences. With QuAPPro, we present the first interactive web app that allows quantification and alignment of polysome profiles, independently of the device or software that was used to generate the profiles. This user-friendly tool does not only speed up the analysis of polysome profiles but also facilitates reproducibility and documentation of the process.

## Introduction

In all organisms and cell types, regulation of protein biosynthesis is crucial for adapting gene expression to environmental cues. After Wettstein *et al*. published the first polysome profiles about 60 years ago (Wettstein et al. 1963), polysome profiling became a widely used method to assess both global and mRNA-specific translation efficiency. This experimental procedure separates mRNAs according to the number of bound ribosomes by sucrose density gradient ultracentrifugation. During elution of the gradient, UV absorbance at ∼ 260 nm is recorded, which mostly reflects RNA content. This profile displays distinct peaks for the small and large ribosomal subunits (40S and 60S in eukaryotes), monosomes and polysomes consisting of increasing numbers of ribosomes (Fig. 1A). More recently, size exclusion chromatography followed by uHPLC was established as an alternative to sucrose density gradients (Yoshikawa et al. 2018). As a common measure for global translation activity, the surface under the polysomal part of the profile (mRNAs with two or more ribosomes) is used as a proxy for the number of ribosomes that are engaged in active translation. In addition, the speed of translation elongation can be assessed in run-off assays where the initiating ribosome is stalled by Harringtonine (Eshraghi et al. 2021; Popper et al. 2024). Defects in ribosome biogenesis specifically affect the 40S and 60S peaks, or can lead to the appearance of additional peaks, so called half-mers, which represent stalled initiating 40S subunits waiting for 60S joining (Li et al. 2009). When fluorescence intensity can be measured in addition to UV absorbance, association of fluorescently tagged proteins with different parts of the polysome profile can be detected (Shaffer and Rollins 2020). Moreover, polysome profiles are used to assess experimental manipulations such as limited enzymatic digest for ribosome profiling experiments (Ingolia et al. 2009; Cenik et al. 2015; Ferguson et al. 2023), and to study polysome assembly in vitro (Kopeina et al. 2008; Afonina et al. 2015). Therefore, the analysis of polysome profiles is an indispensable tool in the field of mRNA translation research.

**Figure 1:**
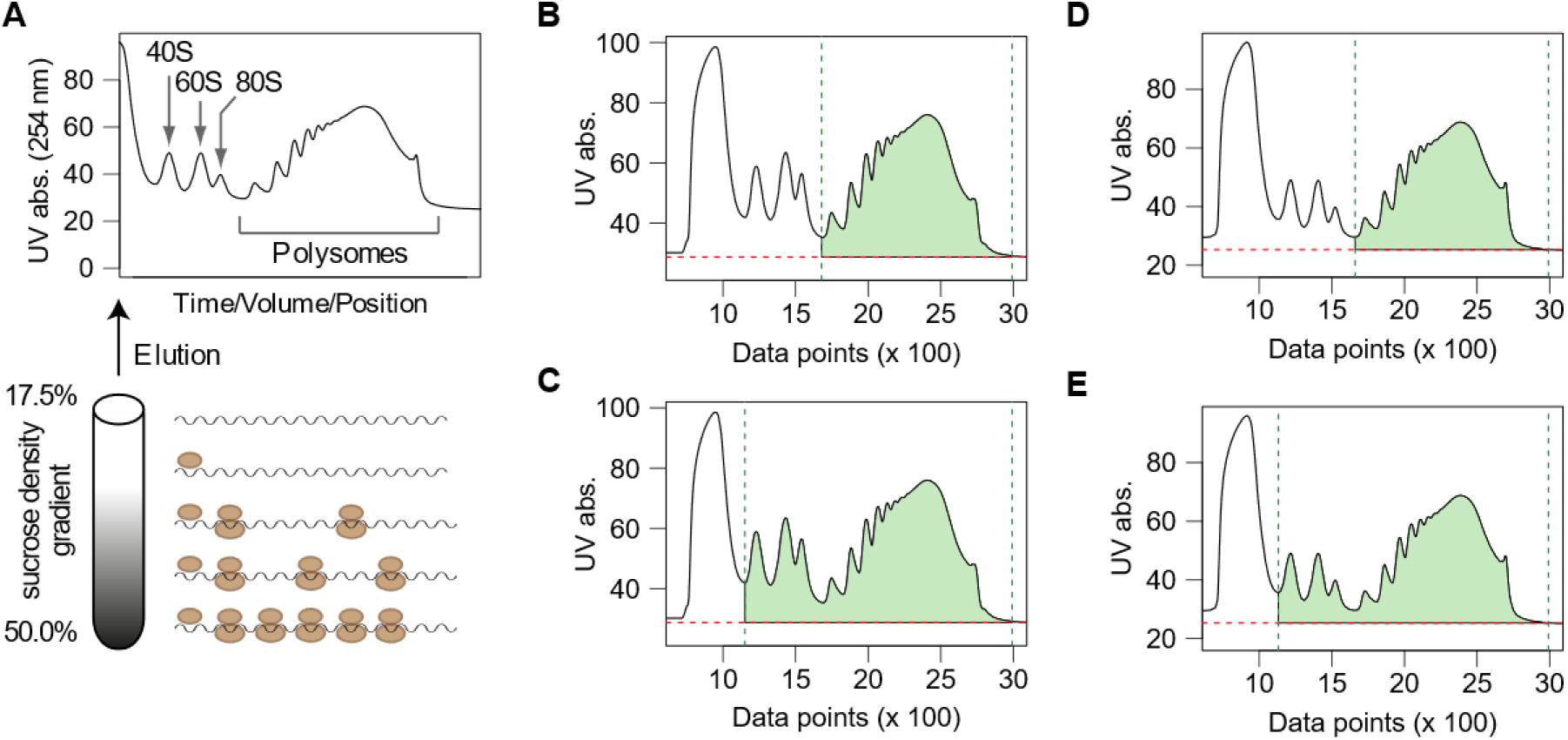
Quantification of polysome profile subsections. **A**. For polysome profiling (here in RAW264.7 cells), mRNAs are separated according to their ribosome load by sucrose density gradient ultracentrifugation. During elution, UV absorbance is recorded. **B**. Polysome profile of RAW264.7 cells with a baseline (dashed red line) and the polysomal part of the profile (green area) as selected interactively by the user. **C**. As in B., with the total ribosomal area in green. **D**. As in B., after 1 h of LPS stimulation. **E**. As in C., after 1 h of LPS stimulation.

With QuAPPro, we present the first interactive web app that allows rapid and convenient alignment of polysome profiles and quantification of profile subsections, independently of the device or software that was used to record the profiles. In addition, corresponding fluorescence profiles can be aligned and quantified for fluorescent polysome profiling.

## Results

### Quantification of polysomal ribosomes

The proportion of actively translating ribosomes is often used as a measure to assess how the overall efficiency of protein biosynthesis is changed under specific treatment conditions. For this purpose, the area under the polysomal part of the profile (with two or more ribosomes; see Fig. 1A) needs to be quantified after subtracting the baseline. For profiles recorded with the Teledyne Isco device (Fig. 1), we suggest to use the UV absorbance at the end of the profile as a baseline, where the gradient has been eluted completely, but sucrose solution still passes the detector. The profiles typically start with a large peak caused by detergents in the lysis buffer (Fig. 1B, Fig. S1), which obscures the simultaneous baseline drop that occurs when the solution reaches the air-filled UV detector (Fig. S1). After loading the profile data into the app, the user can interactively select the baseline and the borders of the area that should be quantified (Fig. 1B). Automatic detection of local minima and maxima supports the selection of borders. For example, the user can click into the valley between the monosome and the disome peak, and the closest local minimum will be detected and selected automatically. Sometimes, peaks overlap so strongly that they will not be separated by a local minimum. In such cases, it is possible to detect an inflection point as a border (Fig. S2). The area under the polysomal part of the profile is then related to the total area under the profile, as quantified from the 40S peak to the end of the polysomes (Fig. 1C). The preceding peak is not included in the quantification, because it mostly reflects components of the lysis buffer (Fig. S1). The results of the quantification can be downloaded as a csv file for further analysis by the user. The profile displayed in Fig. 1B and C was obtained from RAW264.7 cells, a mouse macrophage cell line. In Fig. 1D and E, RAW264.7 cells were stimulated for 1 h with lipopolysaccharides (LPS), which induces the production of numerous inflammatory mediators. Short-term LPS treatment leads to a small but highly reproducible increase of polysomal ribosomes (Schott et al. 2014) and was used here to showcase the ability of our tool to detect these changes. Quantification with QuAPPro reveals that polysomes (Fig. 1B) encompass 75% of the total area (Fig. 1C) in unstimulated cells, while LPS stimulation leads to a small increase in polysomal ribosomes to 81%.

### Alignment and Normalization

The effect of LPS-stimulation on polysomal ribosomes can be further visualized when profiles from the control and treatment conditions are aligned and displayed in one plot. In addition to the baseline, the user needs to select a point on the x-axis along which the profiles should be aligned (the so-called “x-anchor”). For the alignment in Fig. 2A, the local minimum between the monosomes and the disomes was selected for each profile. The small increase in polysomal ribosomes induced by LPS-treatment, however, is difficult to discern with the alignment in Figure 2A, because different amounts of material were loaded onto the two gradients. Therefore, QuAPPro offers the possibility to normalize for the total area under the profile, as quantified in Fig. 1C and E for the control and the treatment condition, respectively. Only when this normalization is applied, it becomes apparent that the polysomal part increases relative to the sub-polysomal section of the profile after the cells were stimulated with LPS (Fig. 2B). Likewise, elution speed or temporal resolution may vary between experiments or instruments. In Fig. 2C, the gradient of the control condition was eluted faster than the gradient of the treatment condition. This difference can be accounted for when also the lengths of the profiles are adjusted according to the lengths of the quantified total areas (Fig. 2D). Figure 2E shows profiles measured after addition of Harringtonine, a compound that selectively stalls initiating 80S ribosomes and therefore allows the measurement of translation elongation speed. Also in this case, normalization to the total area helps to visualize the systematic decrease of polysomes relative to monosomes (Fig. 2F).

**Figure 2:**
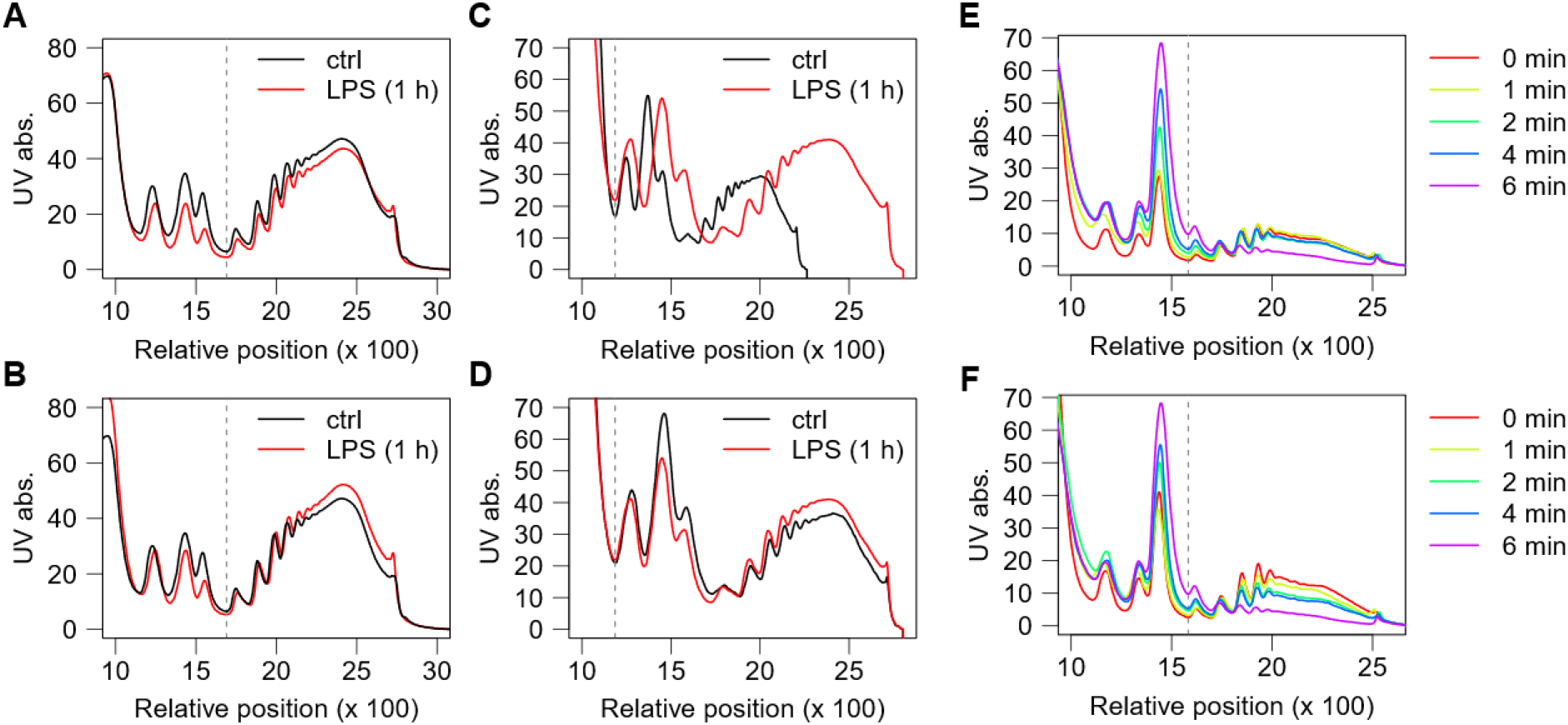
Alignment of multiple profiles along baseline and x-anchor. **A**. Alignment of the profiles from Fig. 1B and D along the baselines and x-anchors (grey dashed line) selected by the user. **B**. As in A., after normalization to the total areas under the profile (quantified in Fig. 1C and E). **C**. Alignment of polysome profiles recorded with different speed from RAW264.7 cells with and without LPS-stimulation. **D**. As in C., after normalization to both surface and length of the total area. **E**. Harringtonine was added to RAW264.7 cells to measure translation elongation speed. To visualize the gradual decrease in polysomal ribosomes, profiles are aligned along the baseline and a position on the x-axis (x-anchor) as selected by the user (grey dashed line). **F**. As in E., after normalization for the total area under the profile.

### Parallel quantification of UV and fluorescence profiles

The ribosome is a large protein complex that interacts with numerous regulatory factors such as initiation or elongation factors and chaperones that assist protein folding. Association of accessory proteins with ribosomal subunits or polysomes is often investigated using Western Blotting on protein samples isolated from fractions of sucrose density gradients. As an alternative, fluorescent polysome profiling with the TRIAX flow cell allows the parallel recording of UV absorbance and the distribution of fluorescently tagged proteins along the gradient (Shaffer and Rollins 2020). Figure 3A shows a fluorescence profile of yeast cells expressing GFP-tagged SBB protein, the yeast homologue of the chaperone Hsp70, which assists in protein folding of nascent peptides (Doring et al. 2017). With QuAPPro, users can select baselines for both UV and fluorescence profiles separately. The area corresponding to the section selected for quantification in the UV profile is then also quantified in the fluorescence profile (Fig. 3A). The fluorescence profile shows several spikes which can be removed in QuAPPro by smoothing the profile (Fig. 3B), which is also possible for UV absorbance profiles, if necessary.

**Figure 3:**
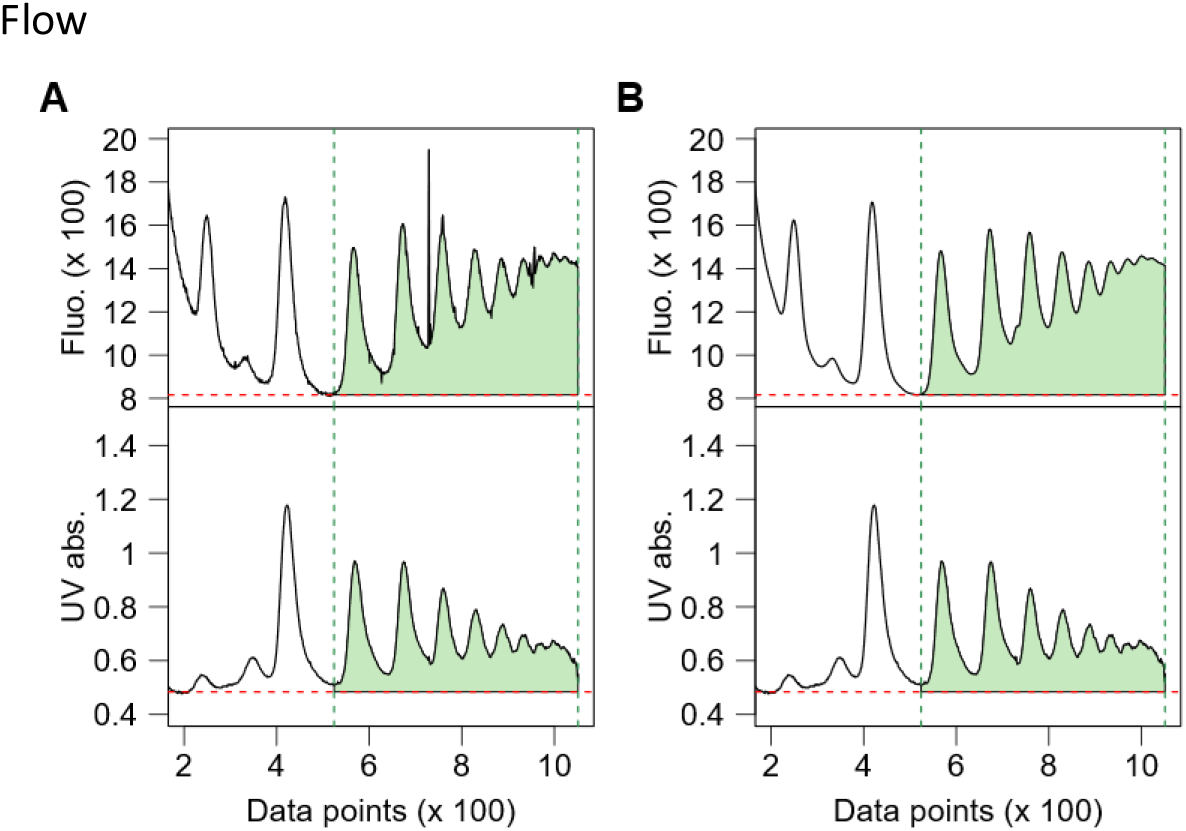
Quantification of fluorescence profiles. **A**. UV and fluorescence profile of yeast cells expressing GFP-tagged SBB protein. The selected region of the UV absorbance profile is quantified in parallel for the fluorescence signal as defined by the indicated borders (green dashed lines) after subtracting the baseline (red dashed line). **B**. As in A., after smoothing the fluorescence profile.

## Discussion

Some devices for recording UV absorbance offer software with additional options for alignment or quantification of the profiles. The TRIAX FlowCell software, for example, offers the possibility to align profiles along selected peaks and to adjust the baseline. In conjunction with the recently available siFractor (siTOOLs Biotech) as an adaptor, ÄKTA liquid chromatography systems can be used for polysome profiling. This opens the possibility to use the Unicorn software for overlaying profiles and quantifying areas under sections of the profile. In addition, most labs that routinely perform polysome profiling have established a way to align and quantify profiles with spreadsheet interface programs like Excel, or with custom scripts in R/Python. Shaffer et al. provide an R script with their protocol for fluorescent polysome profiling that allows quantification of areas under the profile (Shaffer and Rollins 2020). Li et al. implemented correlation-optimized warping (COW) in MATLAB to align and analyze a large number of polysome profiles for investigating ribosome biogenesis in yeast (Li et al. 2009). In contrast to the COW algorithm, QuAPPro can only stretch or shrink a profile by a given factor. Therefore, differences in elution time or temporal resolution between devices can be adjusted for, while variations in sucrose density may still distort the profiles. The accuracy of quantification is affected when neighboring peaks overlap strongly. In some cases, a deconvolution of the profile into its underlying peaks might be necessary for a more precise quantification, for example when monosome peaks need to be quantified that overlap strongly with the 60S peak (Fig. S2). This could be attempted by commercial tools like the peak fitting tool of OriginPro. Despite these limitations, QuAPPro combines several advantages of the approaches mentioned above. No programming skills are required to use QuAPPro. The app allows users to import many different text file formats as input, so that it is not limited to data of a specific device or software. Due to the functions for normalizing the total area and length of the profile, even profiles recorded with different devices can be aligned and analyzed together. It is interactive and therefore highly flexible, because users can decide along which peaks or valleys they would like to align or quantify their profiles. Furthermore, QuAPPro offers automatic detection of local minima, maxima, and inflection points, which renders the analysis very efficient, accurate and reproducible. Not only the results of a quantification or alignment can be downloaded. Also, an entire analysis can be exported and re-imported, so that every step of the analysis is well documented. In addition, all plots displayed in QuAPPro can be downloaded as PDF files at any point of the analysis. As a showcase, most graphs presented in the figures above were retrieved from QuAPPro without any further modifications. Therefore, the app does not only provide a fast, robust, and interactive way to quantify polysome profiles, but also generates ready-to-publish plots.

## Materials and Methods

### Polysome profiling experiments

Details concerning culture and treatment conditions, lysis procedure, sucrose density gradient ultracentrifugation and UV or fluorescence profile recording are listed in Table S1. In brief, mammalian cells were lysed with the indicated lysis buffer in the presence of cycloheximide (CHX). After tumbling the lysates for 10 min at 4°C, nuclei and cell debris were removed by centrifugation at 9,300 × g for 10 min at 4°C.

Growing yeast cultures (OD600 of 0.5-0.6 in YPD) were filtered and lysed by mixer milling (2 min, 30 Hz, MM400 Retsch) with liquid nitrogen in lysis buffer (Table S1). The frozen lysate powder was thawed on ice and spun for 2 min at 30,000 g and 4°C to remove nuclei and cell debris.

The lysates were loaded onto sucrose density gradients. After ultracentrifugation at 35,000 – 40,000 rpm for 2 – 2.5 h, UV absorbance was recorded at 254 nm or 260 nm.

### Implementation in R

QuAPPro was written in R (R Core Team 2021), using the following additional packages: Shiny for building an interactive web app (Chang et al. 2023), shinythemes for the overall appearance of the Shiny application (Chang 2021), colourpicker for interactively selecting colors (Attali 2023), colorspace for color palettes (Zeileis et al. 2009) and stringr for modifying character strings (Wickham 2022). Further details concerning R functions are described below.

### Input file formats

Profiles generated with the Teledyne Isco device Foxy Jr. were recorded using the PeakTrak software and exported as a pks file. These files contain a one-line header, columns separated by a whitespace, UV absorbance in the third column and a comma as decimal separator (for German system settings). For profiles recorded with the BioComp TRIAX system, data was exported as a csv file from the FlowCell software. If the files include a fluorescence signal, the first 51 lines are skipped, and the 52^nd^ line contains the column headers of the recorded data. The third column provides the fluorescence signal, the fifth column the UV absorbance. Columns are separated by a comma, and a point is used as the decimal separator. If no fluorescence signal was recorded, the first 47 lines of the FlowCell file are skipped, and the UV absorbance is extracted from the fourth column. In addition to the provided import options, the user can adjust the number of lines to be skipped from the top of the file, the decimal and column separator, and the column number of UV and, if applicable, fluorescence signal. For example, also tab-separated files or csv files with a comma as decimal separator and a semi-colon as column separator can be imported. These parameters are passed to the R function read.delim().

### Smoothing of profiles

If smoothing of UV absorbance or fluorescence profiles is necessary, an extension of polynomial regression (spline smoothing) is applied using the R function smooth.spline(). The value selected by the user is passed to the smooth.spline() function as “spar” parameter, unless the value is zero or no value was selected and no smoothing is performed.

### Quantification of areas

For quantification of area sections under the profile, the user interactively selects a baseline and the borders of the area of interest. For the baseline, the data point closest to the user’s click in the vertical direction is used. When the user chooses start and end of the quantified area, QuAPPro provides functions to select the closest local minimum, maximum or inflection point. These functions first smooth the profile with smooth.spline(). A local minimum is then defined as a point or plateau where at least five consecutive preceding values show a decrease and five consecutive following values show an increase, while a local maximum is located between an increase followed by a decrease according to the same criteria. Inflection points are identified as local minima or maxima of the first derivative (represented by the difference between the individual data values). The area under the profile is approximated by the sum of all data points from the selected start to the end position after subtraction of the baseline.

### Alignment of multiple profiles

When several profiles are aligned in the same plot, the user needs to choose both baseline and the so-called “x-anchor”, for alignment along the y- and x-axis, respectively. The x-anchor is selected interactively and can be supported by automatic detection of local minima, maxima or inflection points, as described for the borders of quantified areas. In addition, the total area of profiles can be adjusted to the same value by normalization, to compensate e.g. for different amounts of cellular material on the gradient, or different sensitivity settings of the UV detector. When this option is applied, the areas quantified as “Total” by the user are compared, with the largest total area as reference. The y-values of the other profiles are adjusted by the respective normalization factor. When the elution speed differs between gradients, also the length of the “Total” areas can be used for normalization of the x-values. In this case, also the y-values are adjusted, so that the area under the profile is not affected by the change in length.

## Code and data availability

The R code for running QuAPPro and all data presented in this work are available in our GitHub repository (https://github.com/johannaschott/QuAPPro). QuAPPro can also be accessed as a web tool (https://www.umm.uni-heidelberg.de/biochemie/shiny/).

## Acknowledgements

We would like to thank Prof. Georg Stoecklin for his support and ideas, and all members of the Stoecklin lab (Medical Faculty Mannheim of Heidelberg University) for testing QuAPPro with their polysome profiles. We are furthermore grateful to Dr. Andreas Bohne-Lang (IT department of the Medical Faculty Mannheim) for his help with our QuAPPro online version and Dr. Dorina Liebers for providing the SSB-GFP yeast polysome profiles.

## References

Afonina ZA, Myasnikov AG, Shirokov VA, Klaholz BP, Spirin AS. 2015. Conformation transitions of eukaryotic polyribosomes during multi-round translation. Nucleic Acids Res 43: 618–628.

Attali D. 2023. colourpicker: A Colour Picker Tool for Shiny and for Selecting Colours in Plots.

Cenik C, Cenik ES, Byeon GW, Grubert F, Candille SI, Spacek D, Alsallakh B, Tilgner H, Araya CL, Tang H et al. 2015. Integrative analysis of RNA, translation, and protein levels reveals distinct regulatory variation across humans. Genome Res 25: 1610–1621.

Chang W. 2021. shinythemes: Themes for Shiny.

Chang W, Cheng J, Allaire J, Sievert C, Schloerke B, Xie Y, Allen J, McPherson J, Dipert A, Borges B. 2023. shiny: Web Application Framework for R.

Doring K, Ahmed N, Riemer T, Suresh HG, Vainshtein Y, Habich M, Riemer J, Mayer MP, O’Brien EP, Kramer G et al. 2017. Profiling Ssb-Nascent Chain Interactions Reveals Principles of Hsp70-Assisted Folding. Cell 170: 298–311 e220.

Eshraghi M, Karunadharma PP, Blin J, Shahani N, Ricci EP, Michel A, Urban NT, Galli N, Sharma M, Ramirez-Jarquin UN et al. 2021. Mutant Huntingtin stalls ribosomes and represses protein synthesis in a cellular model of Huntington disease. Nat Commun 12: 1461.

Ferguson L, Upton HE, Pimentel SC, Mok A, Lareau LF, Collins K, Ingolia NT. 2023. Streamlined and sensitive mono- and di-ribosome profiling in yeast and human cells. Nat Methods 20: 1704–1715.

Ingolia NT, Ghaemmaghami S, Newman JR, Weissman JS. 2009. Genome-wide analysis in vivo of translation with nucleotide resolution using ribosome profiling. Science 324: 218–223.

Kopeina GS, Afonina ZA, Gromova KV, Shirokov VA, Vasiliev VD, Spirin AS. 2008. Step-wise formation of eukaryotic double-row polyribosomes and circular translation of polysomal mRNA. Nucleic Acids Res 36: 2476–2488.

Li Z, Lee I, Moradi E, Hung NJ, Johnson AW, Marcotte EM. 2009. Rational extension of the ribosome biogenesis pathway using network-guided genetics. PLoS Biol 7: e1000213.

Popper B, Burkle M, Ciccopiedi G, Marchioretto M, Forne I, Imhof A, Straub T, Viero G, Gotz M, Schieweck R. 2024. Ribosome inactivation regulates translation elongation in neurons. J Biol Chem 300: 105648.

R Core Team. 2021. R: A Language and Environment for Statistical Computing. R Foundation for Statistical Computing, Vienna, Austria.

Schott J, Reitter S, Philipp J, Haneke K, Schafer H, Stoecklin G. 2014. Translational regulation of specific mRNAs controls feedback inhibition and survival during macrophage activation. PLoS Genet 10: e1004368.

Shaffer D, Rollins JA. 2020. Fluorescent Polysome Profiling in Caenorhabditis elegans. Bio Protoc 10: e3742.

Wettstein FO, Staehelin T, Noll H. 1963. Ribosomal aggregate engaged in protein synthesis: characterization of the ergosome. Nature 197: 430–435.

Wickham H. 2022. stringr: Simple, Consistent Wrappers for Common String Operations.

Yoshikawa H, Larance M, Harney DJ, Sundaramoorthy R, Ly T, Owen-Hughes T, Lamond AI. 2018. Efficient analysis of mammalian polysomes in cells and tissues using Ribo Mega-SEC. Elife 7.

Zeileis A, Hornik K, Murrell P. 2009. Escaping RGBland: Selecting colors for statistical graphics. Comput Stat Data An 53: 3259–3270.

