## Supplementary material for "QuAPPro: An R/shiny app for Quantification and Alignment of Polysome Profiles"

### Supplement

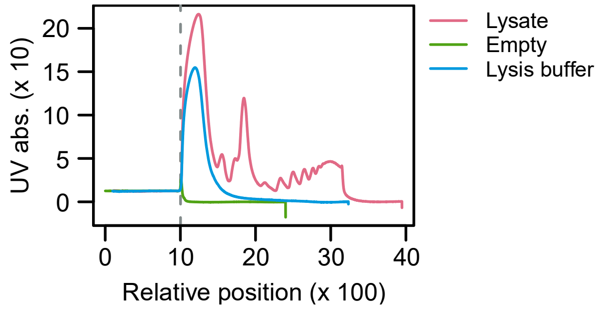


**Figure S1: Baseline of polysome profiles.** UV absorbance was recorded from a sucrose density gradient alone (green line), a gradient loaded with lysis buffer (blue line) or with a lysate of HEK293 cells (red line).

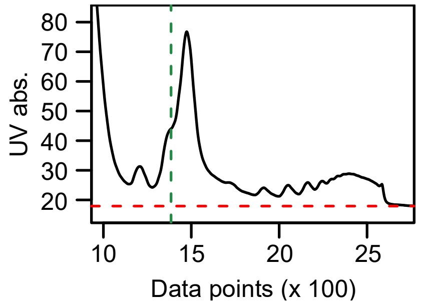


**Figure S2: Detection of inflection points between overlapping peaks.** Polysome profile of mouse embryonic stem cells with baseline (red dashed line) and the inflection point between 60S and 80S (green dashed line) as selected by the user.

| |
|---|
| **Figures 1, 2A – D** |
| Cell line: RAW264.7 |
| Treatment: Stimulated with 100 ng/ml LPS (E. coli O111:B4, Sigma L2630) for 1 h |
| Lysis buffer: 15 mM Tris HCl (pH 7.4), 15 mM $MgCl_2$, 300 mM NaCl, 100 µg/ml CHX, 1% Triton-X-100, 0.1% β-mercaptoethanol, 200 U/ml RNAsin (Promega), 1 complete Mini Protease Inhibitor Tablet (Roche) per 10 ml |
| Sucrose density gradients: 17.5% - 50% (w/v) sucrose (in 15 mM Tris HCl at pH 7.4, 15 mM $MgCl_2$, 300 mM NaCl) |
| Centrifugation settings: 2.5 h, 35,000 rpm at 4°C in a SW60 Ti rotor (Hitachi) |
| Device and software: Teledyne Isco Foxy Jr., PeakTrak software |
| **Figure 2E – F** |
| Cell line: RAW264.7 |
| Treatment: 2 µg/ml HT (Bertin Pharma CAY-15361) for the indicated time periods at RT before addition of 100 µg/ml CHX |
| Lysis buffer: 20 mM Tris HCl (pH 7.5), 150 mM NaCl, 5 mM $MgCl_2$, 1 mM DTT, 100 µg/ml CHX, 1% Triton X-100, 40 U/µl RNasin, 1 complete Mini Protease Inhibitor Tablet per 10 ml |
| Sucrose density gradients: 17.5% - 50% (w/v) sucrose (in in 20 mM Tris HCl pH 7.5, 150 mM NaCl, 5 mM $MgCl_2$) |
| Centrifugation settings: 2 h, 40,000 rpm at 4°C in a SW60 Ti rotor |
| Device and software: Teledyne Isco Foxy Jr., PeakTrak software |
| **Figure 3** |
| Cell line: Yeast BY4741 Ssb1-GFP |
| Lysis buffer: 20 mM HEPES pH 8.0, 140 mM KCl, 10 mM $MgCl_2$, 0.1% NP-40, 100 µg/ml CHX, 1 mM PMSF, 2 × protease inhibitors (Complete EDTA-free, Roche), 0.02 U/µl DNaseI (recombinant DNaseI, Roche), 20 µg/mL leupeptin, 20 µg/mL aprotinin, 1 µg/uL E-64, 40 µg/mL bestatin. |
| Sucrose density gradients: 5% - 45% (w/v) sucrose gradients (in 20 mM HEPES pH 8.0, 140 mM KCl, 10 mM MgCl2, 0.1 mg/ml CHX) |
| Centrifugation settings: 2.5 h, 35,000 rpm at 4°C in a SW40 rotor |
| Device and software: Piston Gradient Fractionator and TRIAX detector (FC-2 dual wavelength flow cell, BioComp), FlowCell software |
| **Figure S1 - S2** |
| Cell lines: HEK293, mESCs |
| Lysis buffer: 20 mM Tris-HCl (pH 7.4), 10 mM $MgCl_2$, 200 mM $KCl_2$, 1% NP-40, 100 µg/ml CHX, 2 mM DTT, 200 U/ml RNAsin, 1 complete Mini Protease Inhibitor Tablet per 10 ml |
| Sucrose density gradients: 17.5% - 50% (w/v) sucrose (in 20 mM Tris-HCl at pH 7.4, 10 mM $MgCl_2$, 200 mM $KCl_2$) |
| Centrifugation settings: 2 h, 40,000 rpm at 4°C in a SW60 Ti rotor |
| Device and software: Teledyne Isco Foxy Jr., PeakTrak software |

**Table S1: Experimental and technical conditions of polysome profiles.** This table lists cell lines, treatment conditions, lysis buffer components, centrifugation settings and information about the devices used for recording the profiles.
